# EPIraction - an atlas of candidate enhancer-gene interactions in human tissues and cell lines

**DOI:** 10.1101/2025.02.18.638885

**Authors:** Ramil N. Nurtdinov, Roderic Guigó

**Affiliations:** Centre for Genomic Regulation (CRG) The Barcelona Institute for Science and Technology, Dr. Aiguader 88, Barcelona 08003, Catalonia, Spain; Universitat Pompeu Fabra (UPF), Barcelona, Catalonia, Spain

## Abstract

Enhancer-gene maps are essential to understand gene regulation. The identification of enhancers contributing to regulating a given gene, however, is challenging as they may lie far from the gene and act in a tissue-specific manner. Here we present EPIraction, an enhancer-gene atlas in the human genome. As H3K27ac is a typical mark of enhancer function, EPIraction relies on an exhaustive collection of 1,538 H3K27ac ChIP-Seq datasets. We used these data together with other epigenomic data to measure enhancer activity. To predict enhancer-gene interactions we first applied the conventional Activity By Contact algorithm to score enhancer-gene pairs in individual tissues. Then, we re-scored the interaction rewarding those present in biologically similar tissues. We used our predictions to annotate the candidate target genes for 1,664,956 SNPs from 3,693 UK Biobank and 760 FinnGen GWAS traits. The EPIraction predictions can be accessed through the portal at https://epiraction.crg.es. The portal can be employed to produce annotations for user-provided lists of SNPs or genomic intervals.

## Introduction

The comprehensive annotation of the human genome is the main goal of the ENCODE Consortium (1, 2). Towards that end, ENCODE has collected thousands of measurements of transcription, high-throughput chromosome conformation capture (Hi-C) contacts, chromatin accessibility, histone modifications, transcription factors binding and other omics assays, identifying millions of functional elements in the human and mouse genomes (2, 3). How these regulatory elements relate to the regulated genes, however, is largely unknown.

Two main approaches exist for predicting enhancer-gene interactions in-silico. The first approach treats the task as a classification or ranking problem (4–6) making tissue-specific predictions from single-tissue data. These algorithms are highly dependent on the quality of input assays and experience difficulties transferring the model from one tissue to another. The second approach uses correlation of some epigenetic and/or transcriptional signals over candidate enhancer-gene pairs across many tissues, samples or individual cells (7–11). These approaches have difficulties in modeling the effect of multiple enhancers in the same gene in a tissue-specific manner and do not utilize the Hi-C data to prioritize distal enhancers.

To overcome the shortcomings of both these approaches, we developed the EPIraction algorithm. We predict enhancer-gene interactions in two steps. Similarly to the conventional ABC algorithm (5) for every enhancer-gene pair we first calculated single-tissue Activity-By-Contact values. In a second step, we recalculated these values, rewarding enhancer-gene pairs in a given tissue that are active in tissues biologically similar to the tissue being scored. We benchmarked our predictions using eQTL associations and ChIA-PET or HiChIP contacts.

Predictions of enhancer-gene interactions are often used to annotate GWAS SNPs (10, 12). GWAS trait-associated SNPs span quite wide regions due to linkage disequilibrium and dissecting the actual causative variant becomes very challenging. Fine-mapping is often used to resolve this problem (13, 14), but the functional context of GWAS SNPs can also be employed (15, 16). We used EPIraction to assign potential target genes to trait-associated SNPs from the UK Biobank (17) and FinnGen (18) and developed a web portal https://epiraction.crg.es that can be dynamically queried with user-provided lists of GWAS SNPs or genomic intervals. The EPIraction predictions are also available through the ENCODE portal https://www.encodeproject.org and UCSC Genome Browser through EPIraction Track Data Hub.

## Results

### The EPIraction algorithm

Similarly to the ABC algorithm we first use epigenomic data to measure enhancer activity and Hi-C data to measure enhancer-promoter contact probability within each tissue and sample. In a first step, we score enhancer-gene pairs in each individual tissue. In the second step, we re-score these predictions based on the predictions observed in biologically similar tissues (see below), rewarding enhancer-gene pairs in a given tissue that are specifically predicted only in the related sets of tissues.

### Candidate enhancer and promoter sets

We used the peaks of open chromatin and CTCF data to produce a set of 1,665,733 tissue-agnostic candidate enhancers and the Gencode v40 annotation (19) to produce a set of 19,283 tissue-agnostic promoters of protein-coding and 12,063 promoters of long noncoding RNA (lncRNA) genes, see Methods. lncRNA genes play a dual role in our analysis: we considered lncRNA promoters as candidate enhancers affecting the expression of nearby genes, as well as genes being regulated by other enhancers (including lncRNAs).

### Activity values

We computed the activity of enhancers as a function of H3K27ac, CTCF, open chromatin (either DNAse-seq or ATAC-seq) and cofactor measurements (either P300 or H3K4me2 or H3K4me1, see Methods). We collected 1,538 H3K27ac ChIP-Seq dataset from ENCODE (3), GEO (20), ENA (21) and BLUEPRINT (22), across 78 tissues, requesting at least ten assays per tissue. Since there is no open chromatin, CTCF and cofactor data for each of the 1,538 H3K27ac samples, to impute these values to candidate enhancers (and to the promoters) in each of the samples, we used the following approach (Fig. 1a, Tables S1 and S2 and Methods). First, we aggregated H3K27ac ChIP-Seq data across all samples from each tissue to produce comparable tissue-consensus H3K27ac values. Next, we used a representative assay from one sample for each of the 78 tissues to compute tissue-consensus open chromatin, CTCF and cofactor values for each candidate enhancer in each tissue.

**Fig. 1.**
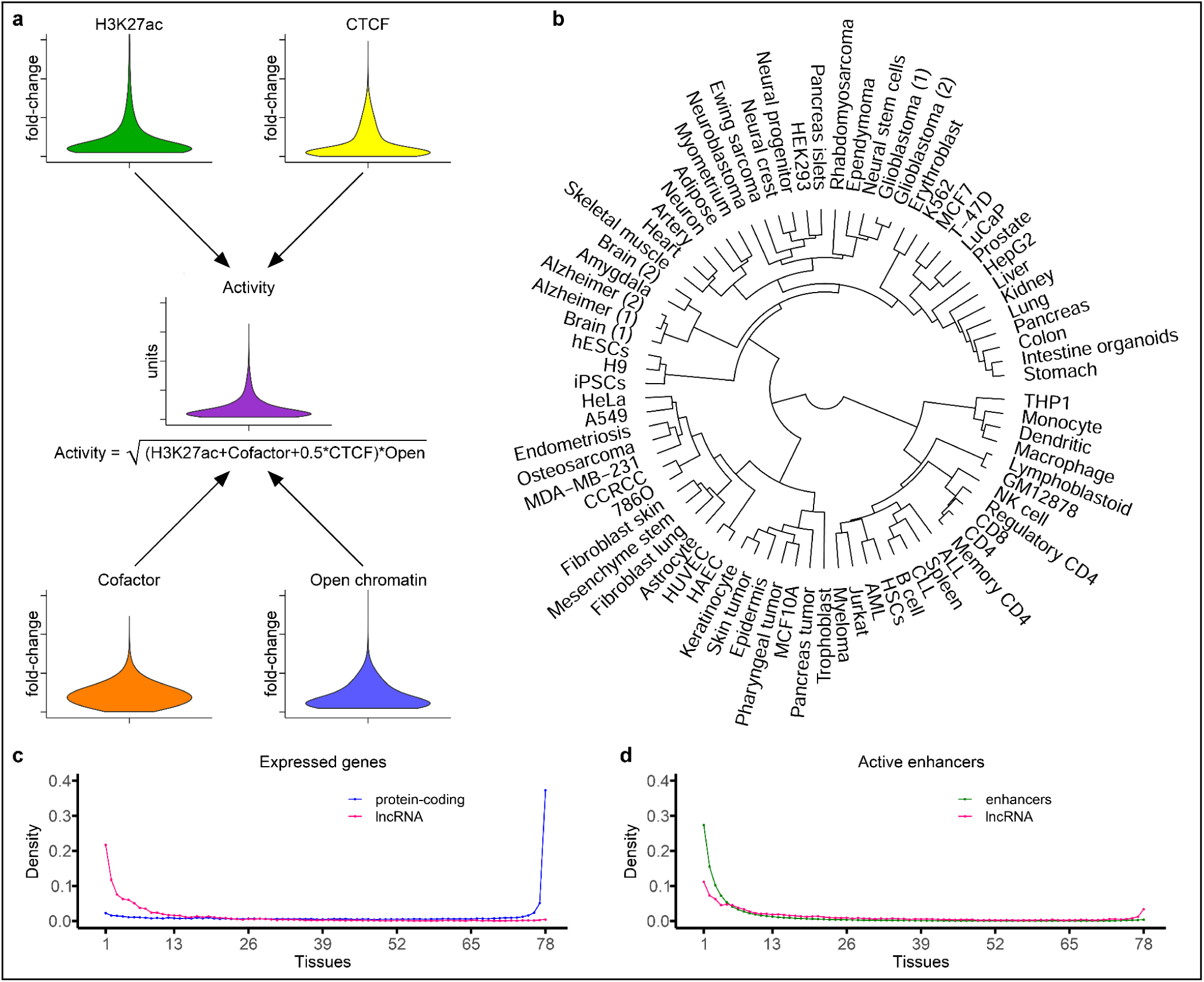
Activity of the enhancers and promoters. **a)** Computing the tissue-consensus and sample-specific activity data for each tissue. Data and plots explain K562 cells. To develop sample-specific activity values we employed H3K27ac from individual samples, to develop tissue-consensus activities we employed tissue-consensus H3K27ac data. **b)** Dendrogram of tissues based on similarity of tissue-consensus activities across promoters and enhancers. **c)** Broad and tissue-specific expression of protein-coding and lncRNA genes. **d)** Broad and tissue-specific activity of lncRNA gene promoter and enhancer regions.

P300, H3K4me2 and H3K4me1 cofactors are well known makers of enhancer activity and shown high correlation with H3K27ac fold-change values on enhancers (Extended Data Fig. 1a-c). Several studies showed synergic effects of CTCF binding and enhancer activities (23–25). We found an inverse relationship between H3K27ac and CTCF signals (Extended Data Fig. 1d), however, for many enhancers, both CTCF and H3K27ac were substantially high: 9,801 (6%) of the candidate enhancers in K562 cells have H3K27ac and CTCF above the value of seven fold-change simultaneously. Considering all the above we calculated the activity for each enhancer region as the geometric mean of open chromatin fold-change and the sum of H3K27ac, cofactor and 0.5 fraction of CTCF fold-change data (Fig. 1a), providing either tissue-consensus H3K27ac values or sample-specific ones.

Fig. 1b shows the clustering of tissues according to the Pearson correlation on tissue-consensus activity of enhancers and promoters. We found a relatively good clustering of tissues: all brain samples cluster together in a dense branch, all primary immune cells cluster together in myeloid and lymphoid branches, cancer samples and cell lines overall lie distantly from the normal tissues.

Protein-coding and lncRNA genes showed different expression patterns. While 6,729 (37%) of protein-coding genes were found expressed with at least 1 TPM in all tissues, only 41 (0.6%) of lncRNA genes are expressed in all our tissues (Fig. 1c). Promoters of lncRNA genes behave similarly to conventional enhancers in terms of activity: approximately 26% of enhancers and 11% of lncRNA promoters are active in one tissue only (Fig. 1d).

### Activity-by-Contact scores

To calculate enhancer-promoter contact probabilities we collected 51 conventional insitu and 28 next generation intact Hi-C data (26) generated within the ENCODE Consortium (2) or available in GEO (20). We quantified Hi-C contact matrices at 2kb resolution and imputed the zero and first diagonals (Methods). We also produced a consensus Hi-C matrix, consisting of approximately 30 billions of intrachromosomal contacts, combining together all intact Hi-C data. We averaged these contacts at different distances to produce the baseline contact map. Eventually, for each enhancer-promoter pair within each tissue we calculated three contact probability values: the one obtained from Hi-C data from this tissue, the one obtained from the consensus Hi-C matrix and the baseline contact probability. The maximum of these three values become the final contact probability (Methods). For each enhancer-promoter pair within each tissue and sample we obtained the Activity-by-Contact (ABC) score as the product of the enhancer activity by the contact probability.

### Conventional ABC predictions

We first applied the conventional ABC algorithm to tissue-consensus data. For each gene, we considered candidate enhancers those laying within 2 Mb from the gene transcription start site (TSS). We summed the ABC scores for promoter and all enhancers and used this value to normalize ABC scores for every enhancer within each tissue. Thus we estimated the contribution of each enhancer and made the total contribution of the promoter and all enhancers equal to one within each tissue.

We investigated how frequently our implementation of ABC algorithm selects tissue-specific enhancers compared with broadly active ones. In total we identified 1,141,734 enhancers that are active in at least one tissue. The ABC algorithm, however, preferentially selects broadly active enhancers: 96.6% of the enhancers that are active in all tissues are in ABC enhancer-gene pairs compared to 13.0% of enhancers active in three tissues and 5.1% of enhancers active in one tissue (Fig. 2a). Two factors are responsible for this bias. Firstly, broad enhancers are, on average, more active (across the tissues where they are active) than tissue-specific enhancers (Fig. 2b). Secondly, broadly active enhancers are located closer to TSS than tissue-specific ones (Extended Data Fig. 2).

**Fig. 2.**
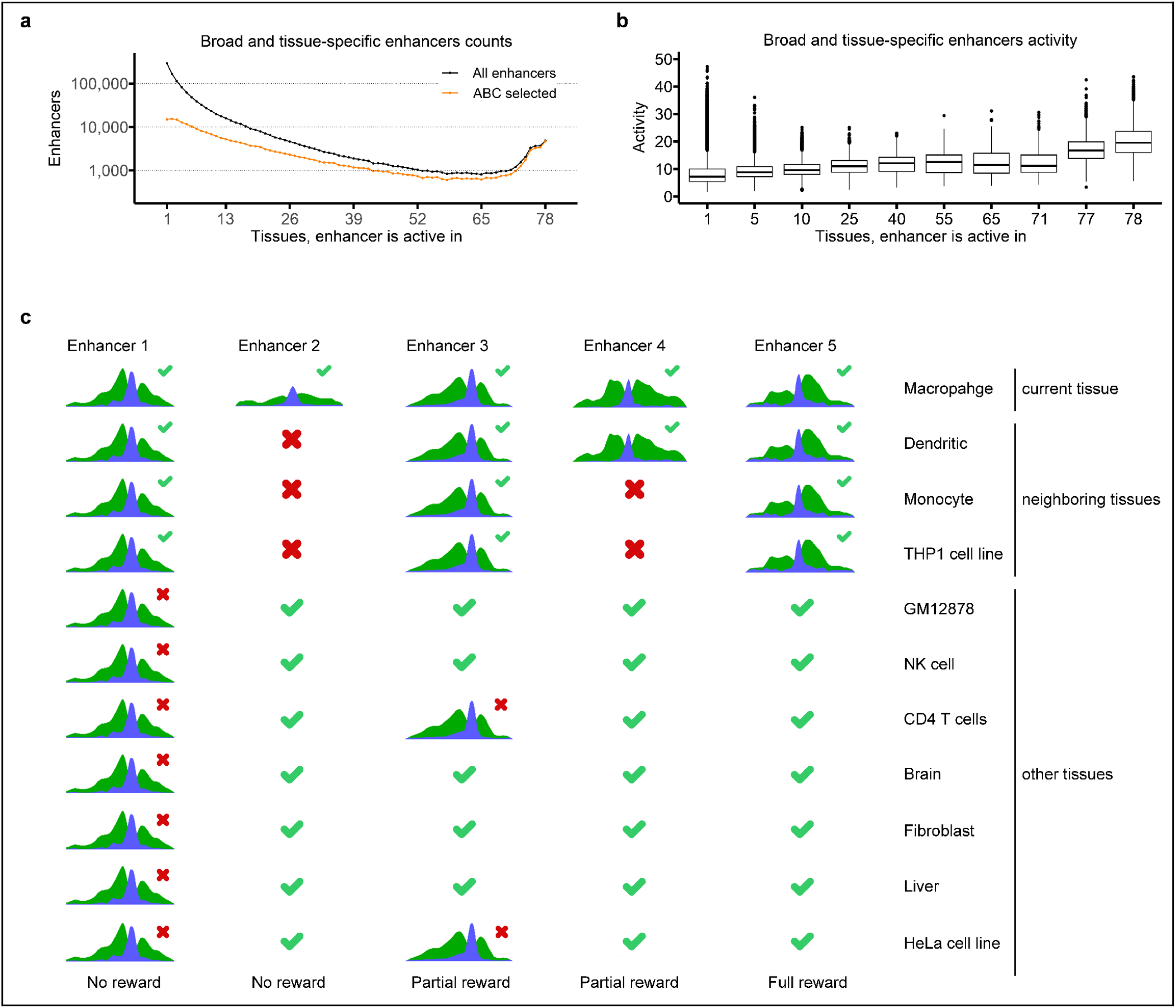
ABC and EPIraction algorithms. **a)** Number of enhancers included in ABC predictions at 0.05 impact threshold and total number of enhancers as a function of the number of tissues in which the enhancer is active. **b)** Activity of enhancers as a function of the number of tissues in which the enhancer is active. **c)** Overview of the second step of the EPIraction algorithm. In this step enhancer-gene pair scores are updated depending on the set of tissues in which the enhancer is active. The green check mark on top of the enhancer indicates that enhancer presence in this tissue contributes favorably to the score. The green check mark alone indicates that enhancer absence is favorable. The red cross on top of the enhancer indicates that enhancer presence in this tissue is disfavorable. The red cross alone indicates that enhancer absence is disfavorable. Enhancer 1 is active in all tissues and thus enhancer-gene pairs containing this enhancer are not rewarded. Enhancer 2 is active only in the target tissue, and enhancer-gene pairs containing this enhancer are not rewarded either. Enhancer 3 is active in all neighboring tissues and some other tissues and thus gains partial reward. Enhancer 4 is active in one of three neighboring tissues and thus gains partial reward. Enhancer 5 is active in all neighboring tissues and absent in all other tissues and thus gains full reward.

### EPIraction predictions

We took advantage of ABC values calculated across many tissues and samples to enhance the contribution of enhancers active in similar tissues. This analysis is based on sample-specific ABC values. For every gene, we produced the Activity-By-Contact matrix. The rows of this matrix correspond to all 1,538 samples, the columns of this matrix correspond to gene promoter and all nearby enhancers. To perform the prediction for the particular gene in the particular tissue, we first defined the neighboring tissues for the given gene in the given tissue. To do so we calculated the average Pearson correlations (row-wise) between the gene’s enhancers activities in samples from this tissue and the samples for each of the other tissues. We considered only samples in which the expression of the gene is greater than 1 TPM. We set a minimum threshold for Pearson correlation to 0.85. The top three tissues with the highest correlation are considered the neighbor tissues for the given gene in the particular tissue.

Then, for every enhancer-gene pair we calculated three ABC scores. The **ABC_tissue** score is the average of the ABC enhancer-gene pair scores across the samples from the tissue in which we are making predictions. The **ABC_neighbor** score corresponds to the average ABC score in samples from the neighboring tissues (if any). The **ABC_complete** score corresponds to the average ABC score across all 1,538 samples. Finally, we re-scored each enhancer-gene prediction by adding the difference between **ABC_neighbor** and **ABC_complete** scores, only if this difference is positive. This rewards specifically the enhancer-gene predictions that are active only in the scoring tissue and in neighboring tissues. Enhancers active only in one or active in all tissues are not rewarded (Fig. 2c). We found that this simple approach substantially corrects the ABC bias towards broadly active enhancers.

### Predicted enhancer-gene pairs

In total we found 18,048 protein-coding and 6,810 lncRNA genes being sufficiently expressed in at least one tissue. On average, across all tissues, protein-coding genes showed higher expression and promoter activities compared to lncRNAs: 78.9 vs and 13.2 TPMs, and 24.8 and 12.4 activity units respectively. At the 0.01 threshold, in total, across all 78 tissues, we predicted 3,194,630 unique enhancer-gene pairs (2,772,901 protein-coding and 421,729 lncRNA). Among these predictions, 1,995,948 pairs were observed in at least two tissues (1,777,514 protein-coding, 218,434 lncRNA), 948,536 pairs were observed in at least five tissues (870,820 protein-coding, 77,716 lncRNA) and 6,345 pairs were observed in all 78 tissues (6,287 protein-coding and 58 lncRNA).

On average per tissue, at the 0.01 threshold, we predicted 253,870 interactions: 234,836 for protein-coding genes and 19,034 for lncRNAs (Table S3). We predicted at least one enhancer for 18,048 protein-coding and 6,809 lncRNA genes. On average, across all tissues, we predicted 153.6 and 61.9 unique enhancers for protein-coding and lncRNA genes respectively. At the 0.05 threshold, on average per tissue, we predicted 40,053 interactions: 36,716 for protein-coding genes and 3,337 for lncRNAs (Table S4). In total across all tissues we predicted 557,461 unique enhancer-gene pairs (482,994 protein coding and 74,467 lncRNAs).

### Benchmark of EPIraction predictions

Properly benchmarking enhancer-gene predictions is challenging as we lack a proper set of known enhancer-gene interactions across many tissues. Here, to benchmark EPIraction, we used the fine-mapped eQTLs from the EBI-EMBL eQTL Catalogue (27). We expect regulatory SNPs mapping to enhancers to be enriched among eQTLs for the predicted target genes. We also used ChIA-PET and HiChIP data which captures physical interactions between different regions of the genome. We expect ‘bona fide’ enhancer-promoter pairs to be enriched in physical contacts. We compared EPIraction enhancer-gene predictions with those by the most recent version of ABC and the comprehensive ENCODE-rE2G predictions (6) and with the EpiMap enhancer-gene interactions (10).

### Enhancer-gene distance

Altogether we obtained 32 tissues for which we have enhancer-gene predictions from all four methods and at least one benchmarking assay (Table S5). EPIraction, ABC and ENCODE-rE2G report approximately 3 millions of enhancer-gene pairs per tissue, while EpiMap reports approximately 195 thousand pairs applying a minimal threshold of 0.714. To make the results comparable across methods when computing the enhancer-gene distances, we set the minimal score thresholds to ENCODE-rE2G, ABC and EPIracition algorithms to obtain, on average, the same number of predictions as EpiMap did. EpiMap preferentially predicts enhancer-gene pairs at short distances, 84 kb on average. ABC predicts the most distant enhancers, the average enhancer-gene distance corresponds to 247 kb. ENCODE-rE2G and EPIraction show an intermediate pattern, 160 and 140 kb respectively (Fig. 3a). We dropped these thresholds and applied different ones for analyses below.

**Fig. 3.**
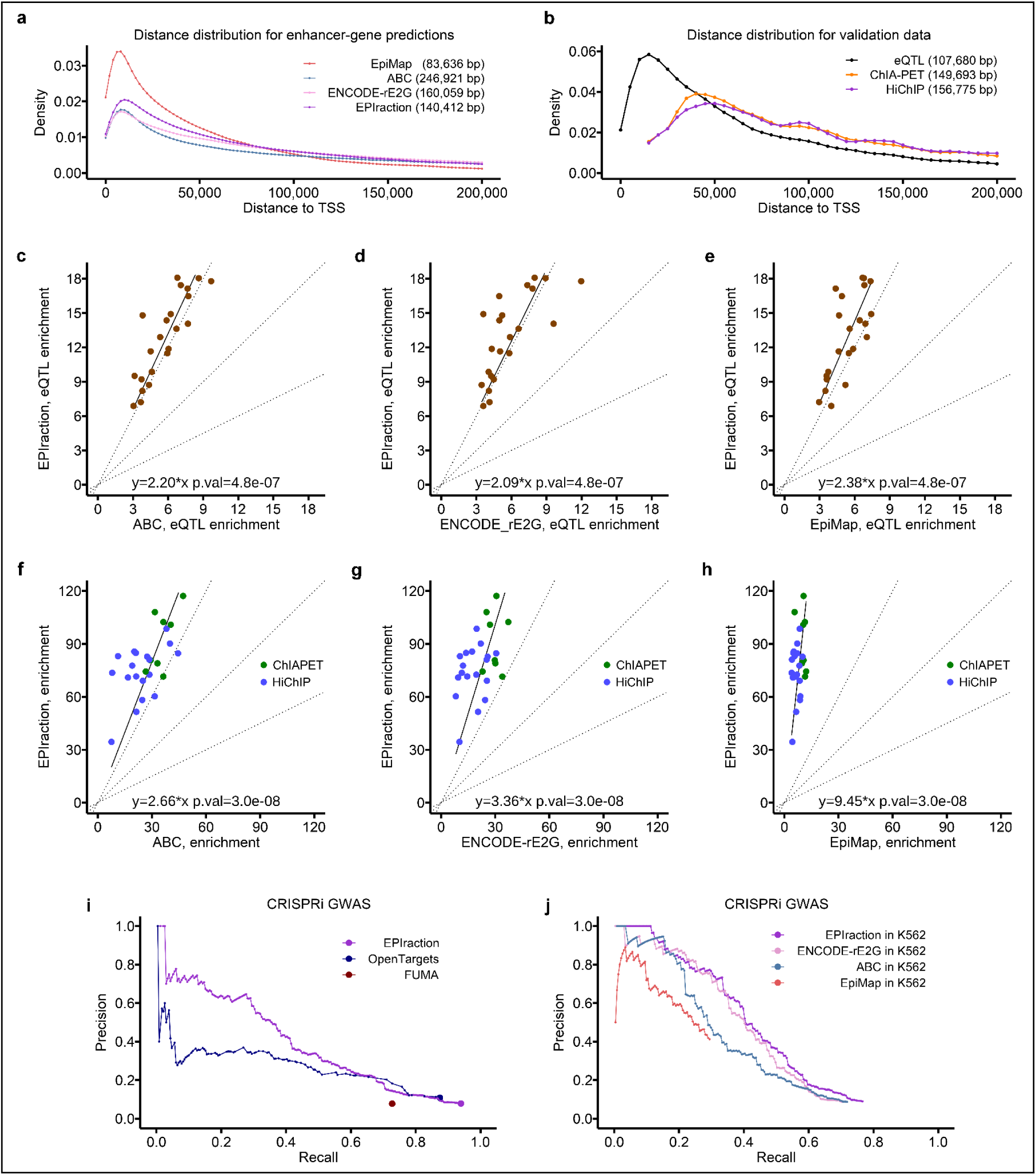
Benchmark of enhancer-gene linking methods. **a)** Distribution of the distance to TSS of EpiMap, ABC, ENCODE-rE2G and EPIraction predictions (32 tissues). Average distance is reported within brackets. **b)** Distribution of the distance to TSS of eQTL and ChIA-PET and HiChIP validation data. **c-e)** Comparative eQTL enrichment in EPIraction enhancer-gene predictions compared to those of other methods of at average recall of 15% across 22 tissues. P-values are calculated by non-parametric paired Wilcoxon test. **f-h)** Comparative ChIA-PET and HiChIP enrichment in EPIraction enhancer-gene predictions at an average recall of 20% across 8 and 18 tissues respectively. Loops are weighted by the contact probabilities. **i)** Comparative performance of EPIraction in tissue-agnostic mode, OpenTargets and FUMA algorithms predicting enhancer-gene contacts at GWAS loci. **j)** Comparative performance of EPIraction, ENCODE-rE2G, ABC and EpiMap predicting enhancer-gene contacts at GWAS loci in K562 cells.

### eQTL benchmark

We collected eQTL data for 23 tissues from EBI-EMBL eQTL Catalogue. The authors use whole genome data for 3,202 individuals (28) to impute SNPs. As a consequence there is a substantial number of eQTLs that are in strong linkage disequilibrium. Analyzing the original whole genome data we found 2% of SNPs lie within promoter regions of protein-coding genes, 18% of SNPs lie in strong (r^2^>0.5) and 35% lie in moderate (r^2^>0.2) linkage disequilibrium with promoter ones. As a consequence, we found approximately 67% of fine-mapped eQTLs either within promoter regions or in strong linkage disequilibrium with promoter SNPs. The remaining 33% of promoter-independent eQTLs are also in substantial linkage disequilibrium in-between. Eventually only 3.6% of fine-mapped eQTLs overlap enhancers, predicted by any of the enhancer-gene linking algorithms, and thus, are suitable for benchmarking purposes (Table S6). The average distance between eQTL and target gene corresponds to 108 kb (Fig. 3b).

Without thresholding EPIraction is able to identify, on average, 45% of eQTL associations, followed by ENCODE-rE2G, ABC and EpiMap with 36%, 34% and 15% respectively. To make the results comparable among methods, we set the score threshold for each algorithm to predict a similar proportion of eQTL associations, that is, to obtain a similar recall. Then, as a metric of performance we computed the enrichment of eQTL associations over expected in the predicted enhancer-gene pairs. As a background, we used the set of common SNPs from dbSNP (29). We computed the eQTL enrichment as the fraction of eQTL associations supported by enhancer-gene predictions over the fraction of common SNPs that map to the enhancers in enhancer-gene predictions. The higher the accuracy of the enhancer-gene pair predictions, the lower the number of predictions required to achieve a given recall; thus, the lower the number of common SNPs mapping to the enhancers, and the higher the enrichment. EPIraction shows the highest enrichment among all methods, followed by ABC, ENCODE-rE2G and Epimap at 15% recall (Fig. 3c-d) and at 30% recall (Extended Data Fig. 3a, b).

### ChIA-PET and HiChIP benchmark

We extracted a list of contacts (significant loops) from the ChIA-PET and HiChIP data, and computed recall as the fraction of contacts supported by at least one enhancer-gene pair prediction (i.e. the enhancer and the gene promoter lie in contacting regions). Similarly to eQTL data we kept the significant contacts that connect promoters of protein-coding genes with enhancers that were predicted by at least one method (Table S7). As the background set to compute the enrichment, we used the set of all common SNPs, as in the eQTL benchmark. EPIraction achieved a maximum average recall of 79%, followed by ENCODE-rE2G with 78%, ABC with 74% and EpiMap with 22%. Similarly to the eQTL benchmark, we compute the enrichment in observed interactions at comparable average recall. We computed enrichment at recalls 20% for all four methods and 50% for all methods without EpiMap (Fig. 3f-h and Extended Data Fig. 3c, d).

### Candidate target gene annotation for non-coding SNPs

The identification of genomic regions involved in the regulation of gene expression can help in the characterization of the molecular mechanisms through which (non-coding) SNPs impact phenotypes. Here, we used EPIraction data to annotate non-coding SNPs from 3,693 GWAS traits from UK Biobank (17) and 760 traits from FinnGen (18) that have at least 100 SNP associations at fdr < 0.1. We overlapped these SNPs with EPIraction enhancers and used the EPIraction enhancer-gene predictions to link GWAS SNPs to target genes in a tissue-agnostic way. In total at 0.01 threshold, across all GWAS traits, we linked 1,664,956 SNPs to 18,036 protein-coding and 6,776 lncRNA genes, for a total of 6,993,734 unique SNP-gene pairs (6,072,990 protein-coding and 920,744 lncRNA). These numbers include SNPs within promoter regions of the target genes. At the 0.05 threshold, we linked 683,744 SNPs to 17,952 protein-coding and 6,687 lncRNA genes, for a total of 1,297,493 unique SNP-gene pairs (1,116,539 protein-coding and 180,954 lncRNA).

We compare the performance of EPIraction with OpenTargets (30) and FUMA (31), two methods that annotate a list of input SNPs, in the ability to predict the target genes for GWAS loci. As a validation set we used the experimental data from Morris and colleagues (32). The authors used a dual-repressor KRAB-dCas9-MeCP2 system to suppress the enhancer activity at fine-mapped blood trait GWAS loci and investigated the expression changes of the surrounding genes in K562 cells. For the fair comparison we ran EPIraction in tissue-agnostic mode instead of K562-specific one. As OpenTargets and EPIraction provide the scores for their predictions, we computed, in these cases, precision/recall curves. For FUMA, we computed one single precision/recall value considering all predictions. EPIraction has a higher AUPRC value than OpenTargets: 0.37 vs. 0.24 respectively. FUMA had a recall of 0.73 and precision of 0.078, since it predicts many target genes at long distances. At this recall, EPIraction had a precision of 0.145 and OpenTargets of 0.182 (Fig. 3i).

Finally we tested how well ABC, ENCODE-rE2G and EpiMap predict these GWAS loci associations. For the fair comparison we selected the predictions made in K562 cells. EPIraction showed the highest AUPRC of 0.424, followed by ENCODE-rE2G, ABC and EpiMap: 0.394, 0.336 and 0.190 values respectively (Fig 3j).

### EPIraction web portal

We built a publicly available database with the EPIraction predictions, which can be downloaded and queried through a portal at https://epiraction.crg.es. We report regulatory SNPs and enhancer-gene pairs genome-wide. Users can provide genomic intervals or gene identifiers. The portal returns, depending on the case, the EPIraction enhancers, the target genes, and associated SNPs, and traits. The users can search specifically for EPIraction enhancers. In this case, the portal returns the levels of H3K27ac, open chromatin, CTCF and cofactor, the activity of the enhancer and the list of candidate target genes in each tissue. EPIraction is also reporting the summary of ChIP-Seq peaks of 1,800 transcription factors from ChIP-Atlas (33) that overlap SNPs and enhancer or promoter regions.

We see the main application of EPIraction portal in annotating the candidate target genes for the list of provided GWAS SNPs or genomic intervals. As an example we report here the analysis performed for hypercholesterolemia trait from FinnGen study. In total, at 0.1 fdr the authors report 12,749 SNPs associated with this disease. We used the EPIraction portal to annotate target genes for these variants. At the 0.05 threshold we associated 1,219 (9.6%) SNPs with 867 genes. Functional analysis of these genes with Metascape (34) showed their involvement in cholesterol and lipid metabolism (Extended Data Fig 4a). The conventional analysis of hypercholesterolemia SNPs with Ensembl Variant Effect Predictor (VEP) (35) reveals 106 (0.8%) SNPs that substantially affect the sequence of 76 protein-coding genes (Extended Data Fig 4b,c). These results indicate the relevant phenotypic impact of regulatory SNPs.

## Discussion

Regulation of gene expression is mediated by the cooperative action of promoters and enhancers. Gene promoters can be relatively well identified, as they act constitutively, and are located in the vicinity of transcription start sites. Moreover, generally, there is only one or a few promoters per gene. The identification of the enhancers that contribute to the regulation of a given gene is, however, much more complex. There are usually many enhancers regulating a given gene, they are located distal to the regulated gene, often at very large genomic distances, and they tend to have a tissue/condition specific activity.

Precisely identifying the enhancers that participate in the regulation of expression of a given gene, and the tissue/condition in which they are active is essential to understand the regulatory programs of genes. Associating enhancers to genes is, therefore, a very active area of research in computational biology and a number of methods have been recently developed towards such aims (5, 8–10) (11).

Here, we report EPIraction, an enhancement of the ABC algorithm that corrects the bias towards broadly active enhancers when predicting enhancer-gene interactions. The method produces predictions that are comparable or more accurate than current state of the art methods. However, EPIraction lags in comprehensiveness and computation costs. For example, to run ENCODE-rE2G only open chromatin data (read alignment and peak files) are required, the remaining data, including tissue-agnostic Hi-C, is obtained from ENCODE. In contrast, to get the maximum benefit of across-tissues predictions, to run Epiraction the full collection of EPIraction data is required, eventually supplementing it with data from new assays, as well as to re-generate the list of tissue-agnostic enhancers. Thus, ENCODE-rE2G provides the most comprehensive list of 1,458 predictions at ENCODE portal, while EPIraction reports only 78 highly accurate predictions.

Reliable enhancer-gene interaction maps, by uncovering genomic regions involved in the regulation of gene expression, help to explain the mechanisms by means of which non-coding variants impact phenotypes at all levels. Indeed, an important application of the enhancer-gene interaction maps is the annotation of GWAS SNPs, which overwhelmingly occur in the non-coding regions of the genome.

Several studies have shown substantial overlap of GWAS loci and enhancer regions: approximately 60% of autoimmune SNPs overlap immune-cell enhancers (36) and 57% of the SNPs in the GWAS catalog (37) overlap DNase I hypersensitive sites from ENCODE tissues (38). Colocalization of GWAS and eQTLs has been widely used to annotate GWAS SNPs with target genes (30, 31, 39). However, a number of recent studies have highlighted the limitation of this approach (40–42). The use of enhancer-gene linking strategies has become an alternative to annotate GWAS SNPs leading to the annotation of tens of thousands non-coding GWAS loci (10, 12, 43). Here, we have used the EPIraction algorithm to link 1,664,956 unique GWAS SNPs to candidate target genes. Remarkably, we have been able to link GWAS SNPs to 6,776 lncRNA genes, via their associated enhancers. While the functional role of lncRNA genes is often questioned, our results suggest that in a non-insignificant fraction of the cases, they may contribute to human phenotypes. Through the EPIraction portal we provide a user-friendly interface to the EPIraction predictions and the transparent framework for annotation of new data.

We believe, therefore, that EPIraction is an important contribution to the existing enhancer-gene interactions maps. These are essential to understand genome function.

## Data availability

Our results are freely available through a web portal https://epiraction.crg.es. The candidate enhancer-gene interactions and information about gene expression and enhancer activities from 78 EPIraction tissues are also available at ENCODE portal https://www.encodeproject.org/search/?type=Annotation&searchTerm=EPIraction. Analysis code is available at https://github.com/guigolab/EPIraction

## Acknowledgements

We thank the members of ENCODE distal regulation working group for useful discussions and for generating a successful collaboration/competition environment. We thank the members of the ENCODE consortium for generating the large amount of primary data needed to develop EPIraction. We thank Emilio Righi for the help in designing the EPIraction web portal and Emilio Palumbo for computation advice. We also want to acknowledge the participants and investigators of the UK Biobank and FinnGen studies. We acknowledge the support of the Spanish Ministry of Economy and Competitiveness (MEC) (BIO2011-26205) and National Institutes of Health (1U24HG009446-01) grants. We acknowledge support of the Spanish Ministry of Science and Innovation through the Centro de Excelencia Severo Ochoa (CEX2020-001049-S, MCIN/AEI /10.13039/501100011033), the Generalitat de Catalunya through the CERCA programme and to the EMBL partnership.

## Author contributions

R.N. conceived the project and performed analyzes, R.G. supervised the project. R.N. and R.G. wrote the manuscript.

## Methods

### Data processing and normalization

#### Processing RNA-seq data

When selecting RNA-seq data for our analysis we gave preference to designs that include polyA RNA selection followed by stranded library preparation and paired-end sequencing. However for a few tissues we used total RNA or not-stranded data or both. We downloaded raw RNA-seq reads from ENCODE (3) or GEO (20) portals. We mapped these reads to the human (hg38) genome and Gencode v40 transcriptome with STAR (44). Initial expression of genes, as TPM (Transcripts Per Million), was quantified with RSEM (45). Some tissues had a lot, up to 32%, transcripts derived from mitochondrial genes. In order to make gene expression comparable we kept only 19,283 protein-coding and 12,063 lncRNA genes (see below) and re-calculated their TPM values. To deal with genes with very high expression levels (tens of thousands TPMs) in some tissues, we limited the expression of any gene to 2,000 TPMs (that is, we set the expression of the gene to 2,000 TPMs, whenever this value was exceeded), and re-calculated TPMs for the rest.

#### Processing ChIP-Seq data

We directly downloaded fold-change over control (hereafter, fold-change) H3K27ac ChIP-Seq data from ENCODE (3) and BLUEPRINT (22) portals. For H3K27ac data from other sources, as well as for H3K4me2, H3K4me1, CTCF and P300 ChIP-Seq we generated fold-change values ourselves. We downloaded the raw read sequences from the original source. We kept only the first read in case of pair-end data. Reads were aligned to the human genome (hg38) with BWA (46). We ran MACS2 (47) in single-end mode to generate genome-wide fold-change data. We requested a constant fragment length of 150 bp (--nomodel --extsize 150) and removed potential PCR duplicates (--keep-dup 1). Finally, we decreased the resolution of all our ChIP-Seq data, by averaging the signal within each 25 bp bin along the human genome (121,241,684 25-bp bins in total).

#### Processing open chromatin data

When selecting open chromatin data, we gave preference to pair-end DNASe-seq data then to pair-end ATAC-seq data. We downloaded the raw read sequences from the corresponding source and aligned them to the human genome (hg38) with BWA (46). We next filtered out alignments with fragment length above 150 bp with Sambamba (48). We ran MACS2 in pair-end mode and generated genome-wide fold-change data and peaks at a relatively high p-value threshold (-p 0.001). Since the open chromatin data do not have input controls, MACS2 builds the controls internally and then calculates fold-change values. Finally, we decreased the resolution of our open chromatin data averaging the fold-change signal in 25 bp bins along the human genome.

#### Processing Hi-C data

We directly downloaded intact tissue-specific Hi-C matrices from the ENCODE portal. While these matrices were generated at very high resolution of up to 10 bp, we used Juicer tools (49) to extract the data for 2 kb resolution and above. We processed other Hi-C data from ENCODE and GEO ourselves. To do so we adopted the Juicer pipeline. In brief, pair-end reads were mapped to the human genome (hg38) with BWA (46). The custom Juicer script “chimeric_blacklist.awk” was used to select the proper pair alignments; these alignments in turn were sorted and duplicates were removed. We kept only intrachromosomal contacts and limited their distance by 30 Mb. Hi-C matrices were generated with Juicer Pre command with standard resolutions up to 2 kb and standard normalization methods. We used this pipeline to process HiChIP data from GEO. To process ChIA-PET data we downloaded the pair-end alignment files from ENCODE portal and applied the recommended custom script “bam2pairs” to generate a list of contacting regions. We generated Hi-C matrices from these contacts with Juicer Pre.

#### Developing the consensus Hi-C matrix from 29 intact Hi-C data

We obtained all intra-chromosomal contact counts at 2 kb resolution from each intact Hi-C matrix at distances below 10Mb and merged them into one array. We applied Juicer Pre to these data and generated the average consensus Hi-C matrix.

#### Normalization of open chromatin data

To efficiently incorporate open chromatin data into our analysis we normalized them. To produce normalized fold-change data, we first averaged the raw signal across 25-bp bins along the human genome, as a part of the smoothing procedure explained above. We merged all the open chromatin data into a matrix, the rows of this matrix correspond to all human genome bins, the columns of this matrix correspond to 73 different samples (several EPIraction tissues may rarely share one single open chromatin assay, for example healthy and cancerous keratinocytes). We next applied quantile normalization (50) to this matrix, split the normalized columns into individual assays and re-generated normalized bigWig files.

#### Joined P300, H3K4me2 and H3K4me1 data normalization

As a cofactor we used either P300 or H3K4me2 or H3K4me1 ChiP-Seq data. All these are known to be associated with enhancer activity. We used different assays for different tissues since there is no common assay of sufficient quality available for all of them. We normalized the cofactor data similarly to open chromatin, generating a genomic bin-to-sample matrix and performing quantile normalization. As a consequence, all the assays would have the same distribution of the signal across the binned human genome.

#### CTCF normalization

We normalized the CTCF data similarly to open chromatin, generating a genomic bin-to-sample matrix and performing quantile normalization.

#### H3K27ac data normalization

In total we collected 1,538 genome-wide H3K27ac samples. They came from different studies, used different anti-H3K27ac antibodies and required substantial normalization. Similarly to the open chromatin and cofactor we averaged H3K27ac fold-change values across all 25 bp intervals of the human genome. It was unfeasible to quantile normalize the whole H3K27ac matrix of 1,538 samples (columns) and 121,241,684 25-bp regions of the human genome (rows). To overcome the computation limits we normalized the H3K27ac matrix in chunks. We split the long chromosomes (1-12 and X) into two arms by the centromeric region and kept the remaining chromosomes as a single full-length chunks. We performed the first round of normalization applying tissue-sensitive smooth quantile normalization (51) across all 1,538 H3K27ac samples within each chunk. For the second round of normalization for each tissue we created a matrix that includes all the samples from this tissue (columns) and all the bins from the human genome (rows) and ran quantile normalization for the first time. We next averaged the columns of the normalized matrix to generate tissue-consensus H3K27ac fold-change values, added them to the tissue matrix as a separate column and applied quantile normalization for the second time. Finally we split normalized values (columns) into individual assays and generated normalized bigWig files for every sample and one tissue-consensus file.

### Tissue-agnostic lists of promoter and enhancer regions

#### Promoter regions

We used Gencode v40 (19) gene annotation to generate a set of promoters. We selected all protein-coding non-mitochondrial and non-chrY genes and removed the ones with “overlapping_locus” tag (readthrough transcripts). For every gene we selected the main transcript with “Ensembl_canonical” or “MANE_Select” tags. The Transcription Start Site (TSS) of such transcripts were assigned to the corresponding genes. We set the promoter region for every gene as an interval of 300 base positions (bp) upstream of TSS and 500 bp downstream of TSS. We initially constructed such promoter regions for 19,283 protein-coding genes. We then added long noncoding RNA (lncRNA) genes to our set. We selected the TSS of the main transcript and removed lncRNA genes with TSS lying within the exons or promoters of any protein-coding genes. Eventually we end up with a set of promoters for 12,063 lncRNA genes.

#### Peaks of open chromatin and CTCF data

As reported, we used MACS2 to call peaks on open chromatin and CTCF data. We decreased the resolution of our data from single-base to 25 bp base. Thus we made the position (in chromosomal coordinates) and the length of every peak multiple of 25. For each peak, we demanded at least 100 bp of its length covered by a 3.5 fold-change signal. We allowed this coverage to be discontinuous and kept the most upstream and the most downstream positions as a peak termini. Within each sample we filtered out the peaks overlapping the centromeric regions and promoters. In total we identified 8,539,042 open chromatin (across 73 samples) and 2,377,511 CTCF peaks (across 57 samples).

#### Candidate enhancer regions

We used the peaks of open chromatin and CTCF as a source to generate candidate enhancers. We pooled all samples together and merged overlapping open chromatin and CTCF peaks. This resulted in 1,341,244 merged peaks. The signal at each position of these merged peaks is the sum of the fold-changes of open chromatin and CTCF signals.

We plot the length distribution of merged peaks at Extended Data Fig. 5. The median merged peak length is 275 bp and the average is 426 bp. We assume the enhancers to be nucleosome free open chromatin regions, but histone modifications in the adjacent nucleosomes play an important role in enhancer definition and function. Thus, we generated enhancer regions from the peaks satisfying the following conditions: 1) all peaks are covered (totally or partially) by at least one enhancer region, 2) the minimum length of an enhancer region is 450 bp, 3) the minimum distance between enhancer regions is 25 bp, 4) at least 100 bp of a given enhancer region are covered by one or more peaks. We use a segmentation algorithm that attempts to maximize the number of 450 bp long enhancers, as we made this the desired length of the enhancer region. The algorithm is inspired by the segmentation algorithm employed to build candidate cis regulatory regions (cCREs, (3)). This algorithm extends short peaks and splits long peaks to generate regulatory regions of a given length. Thus, in our case, we attempted to extend short peaks (<450 bp, 920,053, 69%) to 450 bp long enhancer region, split longer peaks (>450 bp, 393,750, 29%) into chains of 450 bp enhancer regions, and kept the peaks that are exactly 450 bp long (27,440, 2%) intact as enhancer regions.

We extended the short peaks in three steps. First, we merge neighboring short peaks if the length of the resulting region does not exceed 450 bp (Extended Data Fig. 6a). Merged peaks 450 bp long were considered candidate enhancer regions. Second, we extended the remaining short peaks (including short merged peaks) to 450 bp. We extended all the peaks simultaneously both upstream and downstream in steps of 25 bp. We stop extending a given peak in one direction if the next iteration would produce an extension closer than 25 bp to the neighbor peak (Extended Data Fig. 6b). Third, we merged the remaining short peaks to adjacent short or long peaks. In this case, the resulting peak would exceed 450 bp, since all shorter peaks were already merged or extended before (Extended Data Fig. 6c). Eventually, all short peaks were either extended and/or merged. In total, we obtained 1,309,652 regions: 904,597 (69%) 450 bp long and 405,055 (31%) longer than 450 bp.

We next attempted to split the long peaks into a chain of 450 bp non-overlapping candidate enhancer regions maximizing the fold-change signal within each region. We started growing the first region from the 25 bp window with the highest signal within the peak, and extended it iteratively in both directions with the neighboring bin with the highest signal until we reached the length of 450 bp. We repeated this procedure while it was possible to generate 450 bp regions within the long peak (Extended Data Fig. 6d).

This iterative procedure frequently keeps substantial gaps between 450 bp regions. To close these gaps we extended them equally to reach a minimum gap of 25 bp (Extended Data Fig. 6e). Frequently, the most up- and down-stream parts of long peaks lie outside from candidate enhancer regions. We created additional 450 bp candidate enhancer regions up- and/or down-stream of the splitted chain of enhancers, if the length of the peak outside candidate enhancer regions is greater or equal than 175 bp (Extended Data Fig. 6f), otherwise we extend the first/last enhancer in the chain (Extended Data Fig. 6g, h).

Eventually, we generated 1,665,733 candidate enhancers: 1,198,405 (72%) 450 bp long and 467,328 (28%) longer than 450 bp. These candidate enhancers cover 804,234,475 bp which constitute approximately 27% of the human genome.

### Prediction of enhancer-gene interaction

#### Activity values

We used the tissue-consensus assays to quantify open chromatin, CTCF, cofactor and H3K27ac signals (maximum fold-change) at promoters and enhancers. We requested two alternative conditions to consider a particular enhancer active in corresponding tissue. The conventional enhancers should have the sufficient levels of H3K27ac and open chromatin signal. The CTCF enhancers should have substantial levels of CTCF signal. Otherwise we set the enhancer activity to zero. For the majority of tissues these minimums correspond to 3.5 fold-change of H3K27ac, 3.5 fold-change of open chromatin and 5.5 fold-change of CTCF. However, for some tissues we increased or decreased these thresholds to achieve a similar amount of active enhancers across all 78 tissues, the exact thresholds are reported within Table S1. We calculated the activity of the enhancer or promoter as a product of tissue-consensus open chromatin, H3K27ac, CTCF and cofactor signals [1]

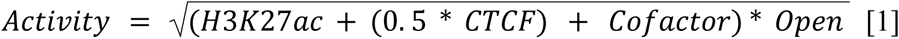

To generate the sample-specific activity values, we applied the same formula [1], replacing tissue-consensus H3K27ac with H3K27ac values from the particular sample.

#### Contact probability from Hi-C data

Hi-C data are stored as sparse matrices binned into equal genomic intervals, the length of these intervals corresponds to the Hi-C matrix resolution. The contacts occurring within the same interval (the zero-diagonal contacts) require special attention since they frequently represent a re-ligation product of adjacent or nearby fragments and thus do not correspond to the valid contacts. Additional attention is required for the contacts that cross the border of two consecutive intervals (the first-diagonal contacts). In principle, they belong to different intervals, but they may actually represent the same re-ligation product. Finally many contacts are supported by only one of very few reads. These contacts are not completely trustful, due to stochastic nature of Hi-C data and/or low mappability of repetitive regions of the human genome (see below).

We quantified Hi-C contact matrices at 2 kb resolution applying SCALE normalization with Juicer tools. Due to the normalization the resulting read counts become fractional. We projected every promoter into a 2 kb interval. If the promoter crosses the border of two intervals, we selected the one with highest overlap. For every promoter we extracted all contacting regions up to 2 Mb. We calculated the contact probability C_n,p_ of the promoter and the distal region as a number of reads that support their contact D_n,p_ divided by the normalization factor M_p_ that is specific for every promoter (Extended Data Fig. 7). We calculated M_p_ as an average value of reads that support two second diagonals in Hi-C matrix [2]. Then, we set the contact probability on the zero diagonal equal to one [3], the contact probabilities on the two first diagonals equal to 0.99 [4], the contact probabilities on the two second diagonals equal to 0.95 [5]. For the third and higher diagonals we set the contact probability as a number of Hi-C reads supporting the contact of the particular region and promoter divided by the M_p_ value [6,7].

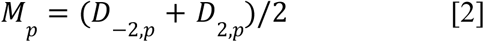

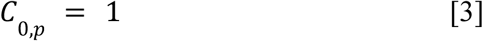

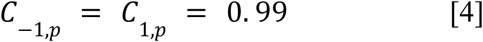

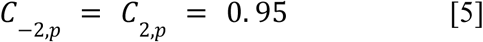

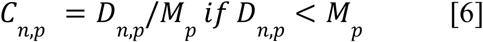

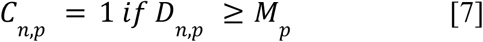

In addition to contact probabilities generated from Hi-C data, we calculated the baseline contact probabilities. To do so we extracted contact probabilities of all promoters with all regions within 2Mb from the consensus Hi-C matrix. To calculate the baseline contact probability on a particular distance, we averaged the values across all pairs at this distance. We used these baseline contact probabilities to constrain Hi-C contacts. If the contact of the promoter and enhancer is supported by one read (before normalization) we constrained their contact probability to the baseline value at the same distance. If the contact is supported by two reads, we constrained the probability to 1.5 the baseline value. If the contact is supported by three and more reads, we did not constrain them.

Eventually, for each promoter-enhancer pair we calculated the contact probability as a maximum value from the corresponding tissue-specific Hi-C matrix or consensus Hi-C matrix or baseline.

#### Generation of the Activity-by-Contact matrix

For each protein-coding and lncRNA gene we generated a single Activity-by-Contact (ABC) matrix from sample-specific enhancer activities and tissue-specific Hi-C contact probabilities. We considered all the enhancers that lie within 2Mb from the corresponding promoter and promoter itself as candidate regulatory regions. The columns of the ABC matrix correspond to all candidate enhancers and to the promoter, the rows of this matrix correspond to all 1,538 H3K27ac samples. The values of this matrix correspond to ABC scores, i.e. enhancer activities in the particular samples multiplied by Hi-C contact probabilities of the promoter and corresponding enhancers in corresponding tissues. The contact probability of the promoter region with itself is considered to be the value of one.

#### EPIraction predictions

We predicted the candidate enhancers for every expressed gene within every tissue independently. When predicting enhancers for a particular gene in a particular tissue, we first removed the enhancers that are inactive in our tissue from the gene ABC matrix. To define the closest tissues we first calculated Pearson correlation of samples from tissue of interest against all other samples within the ABC matrix and then averaged the corresponding correlation coefficients across the samples from every tissue. We defined the closest tissues as the ones with correlation above 0.85 and gene expression above 1 TPM. If we found more than three such tissues we selected the three with the highest correlation.

Then, for every enhancer-gene pair we calculated three ABC scores. The **ABC_tissue** scores are the average of the ABC matrix across the samples from our tissue, the **ABC_complete** scores are the averages across all samples. Finally, if we found at least one closest tissue, we calculated **ABC_closest** scores by averaging ABC scores across the samples from the closest tissues. Based on these scores we calculated the **Reward** for every enhancer-gene pair [6-8].

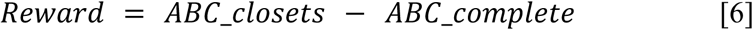

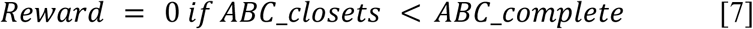

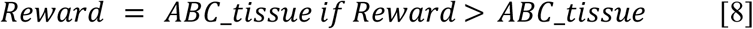

To calculate the contribution of each enhancer we added **Reward** to **ABC_tissue** scores and normalized these values to the sum of **ABC_tissue**. We did not include **Reward** into the normalization denominator.

### Benchmark of EPIraction predictions

We downloaded from the ENCODE portal the ABC (the complete versions when available, for four tissues we used the DNAse-only version) and ENCODE-rE2G predictions, provided by the ENCODE distal regulation group. These predictions were generated for the current hg38 version of the human genome. We extracted the predictions for protein-coding genes and kept the promoter-enhancer pairs at distances below 2Mb. We used the resulting predictions as-it-is, without any modifications or adjustments. We downloaded EpiMap predictions from the EpiMap portal (http://compbio2.mit.edu/epimap_vis). We lifted these predictions from the hg19 to the hg38 genome assembly and extracted the data for protein-coding genes at promoter-enhancer distances below 2Mb. The average length of EpiMap enhancers is 230 bp, the minimum enhancer length is 20 bp. Thus, we extended EpiMap enhancers to a minimum length of 500 nt to be comparable with other predictions.

ABC, ENCODE-rE2G and EpiMap algorithms make their predictions based on individual ENCODE datasets. Therefore, for example, having two DNAse-seq and two H3K27ac datasets for the same tissue they produce four different predictions. In these cases we selected the predictions with the highest number of reported enhancer-gene pairs for the subsequent analysis. In total we created a list of 32 tissues for which we have enhancer-gene predictions from all algorithms and at least one benchmarking assay, Table S5.

#### Common set of enhancers

For every tissue within the validation data we extracted all the enhancers predicted by each algorithm and merged them. We did not merge these sets across tissues, keeping them tissue-specific.

#### Processing of the full genome SNP data

We downloaded the whole genome VCF data for 3,202 individuals (28) from EMBL/EBI portal (http://ftp.1000genomes.ebi.ac.uk/vol1/ftp/data_collections). We converted these files into Plnk2 (52) format, and calculated pairwise linkage disequilibrium statistics with “--r2-unphased allow-ambiguous-allele --ld-window-kb 1000” command.

#### eQTL benchmark

We downloaded fine-mapped eQTL data from the EBI-EMBL eQTL Catalogue (27) portal (http://ftp.ebi.ac.uk/pub/databases/spot/eQTL). We classified a given eQTL as a promoter one if it fell within the promoter region of the protein-coding gene. If the promoter eQTL was found regulating another gene we still considered it as a promoter one, since these genes can interact in *trans*. Similarly, we call all eQTLs that are in sufficient linkage disequilibrium (r^2^ > 0.5) with promoter SNPs as promoter-associated. We considered the remaining eQTLs as promoter-independent and overlap them with enhancers from the same tissue predicted by at least one enhancer-gene linking algorithm (Table S6).

As a metric of performance we computed fold-enrichment of eQTL associations over expected in the predicted enhancer-gene pairs. We considered a particular eQTL supported by an enhancer-gene prediction if the eQTL SNP falls within an enhancer predicted to interact with the eQTL target gene. The enrichment is calculated as the fraction of eQTLs that are supported by enhancer-gene prediction (recall) divided by the fraction of all SNPs in the background that overlap the enhancers that participated in enhancer-gene predictions. As a background we used all common SNPs from dbSNP build 156 (29).

#### ChIA-PET and HiChIP benchmark

We used Hi-C loops (enriched contacts on Hi-C contact matrix) to benchmark enhancer-gene predictions. Similarly to eQTL data, these loops link the gene (the gene promoter region in our case) and the distal genomic loci. We generated Hi-C contact matrices for each ChIA-PET and HiChIP validation dataset above. Although these datasets contain hundreds of millions of contacts, these data are not enough to search for enriched contacts (loops) at small resolutions. We used the 5 kb resolution matrices for loop-calling and limited the distance between contacting regions by 2Mb.

We used three complementary methods to define Hi-C loops: HiCCUPS (49), Peakachu (53) and HiCExplorer (54). HiCCUPS directly uses Hi-C matrices in Juicer format. To run HiCExplorer and Peakachu we performed format conversion. We used cooler software (55) and converted Hi-C matrices in Juicer format at 5 kb resolution and SCALE normalization into Hi-C matrices in HDF5 format. We next rescaled these Hi-C matrices to obtain 250 million contacts and asked HiCExplorer and Peakachu to predict Hi-C loops from these matrices at default parameters. Eventually, we selected predictions that are supported by at least two algorithms.

Similarly to eQTL data we kept only ChIA-PET and HiChIP loops linking promoters and enhancers from the same tissue. As a metric of performance, we computed fold-enrichment of ChIA-PET and HiChIP loops over expected in the predicted enhancer-gene pairs. We considered the particular loops supported by an enhancer-gene prediction if one end overlaps the enhancer and another end overlaps the promoter of the gene predicted to interact with this enhancer. To count ChIA-PET and HiChIP loops, we weighted them by the contact probability. The enrichment is calculated as the weighted fraction of loops that are supported by enhancer-gene prediction divided by the background estimations. Since there are no proper background estimations for ChIA-PET and HiChIP loops we adopted it from eQTL analysis as a fraction of all common SNPs that overlap the enhancers that participated in enhancer-gene predictions.

#### Processing the UK Biobank GWAS data

We used UK BioBank GWAS data linking 11,908 phenotype measurements to polymorphism data in 361,194 individuals (17). Of these, we selected the 4,278 phenotypes common to both sexes. We kept only the phenotypes with quantitative measurements and filtered out phenotypes based on inverted ranks (“continuous_irnt” traits). We applied Benjamini, Hochberg and Yekutieli (fdr) correction to the reported association and kept the ones with fdr<0.1. Eventually, we kept 3,693 GWAS traits with at least 100 fdr-significant associations. We used CrossMap (56) to lift corresponding polymorphisms from hg19 to the hg38 genome assembly. We next used bcftools (57) to map these variants to dbSNP build 156 (29) in order to have a consistent list of identifiers. To annotate the candidate target genes for these variants we overlap them with EPIraction enhancers and report the genes that participate in EPIraction enhancer-gene links at 0.01 score as a candidate target genes.

#### Processing the FinnGen GWAS data

We downloaded the summary statistics for the 2,408 FinnGen GWAS traits from the Google cloud. For only 760 of them we found at least 100 SNPs with fdr < 0.1. Similarly to UK Biobank data we overlap these SNPs with EPIraction enhancers to predict the candidate target genes.

#### Benchmarking GWAS annotation

We used the experimental data from Morris and colleagues (32) to benchmark the performance of GWAS annotation algorithms. The authors used a dual-repressor KRAB-dCas9-MeCP2 system to suppress the enhancer activity at fine-mapped blood trait GWAS loci and investigated the expression changes of the surrounding genes. The authors reported results of 8,192 observations made in two replicates. We considered gene-loci pairs with reported p.value < 0.01 as interacting (positive) and pairs with p.value > 0.5 as non-interacting (negative) observations. We used CrossMap (56) to lift corresponding genomic positions from hg19 to the hg38 genome assembly. The dual-repressor KRAB-dCas9-MeCP2 system efficiently represses the promoter regions at distances up to 1,500 bp away from gRNA location. To efficiently isolate the effect on enhancers, we additionally filtered out the GWAS variants that lie closer than 2 kb from the TSSs. Eventually we end up with 231 positive and 2,933 negative observations.

We downloaded raw OpenTargets data (v22.09) from the EMBL-EBI FTP portal (https://ftp.ebi.ac.uk/pub/databases/opentargets/genetics/22.09). Following the guidelines we built the “v2g_score_by_overall” table under the ClickHouse database engine from these data. We obtained the candidate target genes for our GAWS variants from this table together with the scores generated by OpenTargets algorithm.

To obtain the FUMA predictions, we submitted the hg19 versions of our GWAS variants to the FUMA portal (https://fuma.ctglab.nl/snp2gene). Since FUMA does not supply each GWAS-gene prediction with corresponding score we were able to calculate only a single pair of Precision-Recall values.

### EPIraction web portal

### Collection of regulatory SNPs

We downloaded the full lists of eQTL associations for 109 tissues from the EBI-EMBL eQTL Catalogue (27) portal. We extracted all eQTL associations with p.value < 1e-6 and combined them with fine-mapped ones with p.value < 1e-5. We mapped the eQTL SNPs to dbSNP build 156 with bcftools in order to have a consistent list of identifiers. In total, across the 109 datasets, we obtained 2,848,713 SNPs regulating 16,793 protein-coding genes. These data from 31,863,287 (5,533,411 unique) SNP-gene pairs. We merged these SNPs with 11,273,313 fdr-significant GWAS SNPs from UK Biobank and FinnGen, see above. In total we obtained a list of 12,008,606 regulatory SNPs, for 2,113,397 variants we have both eQTL and GWAS data.

#### ChIP-Atlas peaks

We downloaded the experiment summary file of the ChIP-Atlas database (33) and filtered it for human (hg38) ChIP-Seq (TFs and others) experiments with a valid human gene name in antibody target field. Thus, we obtained a list of 29,708 experiments for 1,800 human transcription factors (17 on average per TF). We downloaded **allPeaks_light.hg38.05.bed.gz** file, which contains the full list of peaks at fdr < 1e-5 (q=5) threshold and filtered it for the peaks from these experiments. We overlapped these peaks with regulatory SNPs, enhancer and promoter regions to make a comprehensive annotation and selected the most significant peak for each TF for the short summary.

#### The portal

We built a publicly available database containing the EPIraction predictions, GWAS and eQTL SNPs and ChIP-Atlas peaks. It can be queried through a portal at https://epiraction.crg.cat. The database contains two types of regulatory information: the regulatory SNPs, derived from UK Biobank, FinnGen and EBI-EMBL eQTL Catalogue, see above, and EPIraction-specific information about enhancer and promoter activities and predicted promoter-enhancer interactions. The “GENOMIC INSIGHTS” and “GENE VARIANTS” tabs in the portal can be used to query the regulatory SNPs and their potential overlap with EPIraction enhancers. The “GWAS ANNOTATION”, “REGIONS” and “ENHANCER-GENE PAIRS” tabs can be used to query EPIraction-specific data. Both these query modes can take as input either a genomic interval or a gene identifier, but provide substantially different results.

The “GENOMIC INSIGHTS” tab of EPIraction portal allows querying the regulatory SNPs. If the user provides a given genomic interval, the portal reports all eQTL and GWAS SNPs within the interval and their associated target genes and traits. EPIraction enhancers and CHIP-Atlas peaks overlapping these variants are also reported. If the user provides a gene identifier, the portal will report the interval data corresponding to region +/– 100 kb from the gene TSS.

The “GENE VARIANTS” tab serves to identify regulatory elements associated with a particular gene. The user provides a gene identifier. The portal reports all eQTL for the gene, all the enhancers that are predicted to regulate this gene in tissue-agnostic mode and all GWAS variants, lying within these enhancers.

The “GWAS ANNOTATION” tab allows querying for candidate target genes given a list of genomic positions or genomic intervals. The user provides a gzipped file that contains a header and a list of genomic positions (for instance, GWAS hits, CpG positions, etc). The portal overlaps these positions with the EPIraction enhancers and promoters, and reports the genes predicted to be regulated by these enhancers, both in a tissue-agnostic and a tissue-specific manner. The md5 hash sum of the input file is used as a job identifier.

The “REGIONS” tab reports specifically EPIraction data. The user provides a given genomic interval. The portal reports all EPIraction enhancers and promoters in that interval. Selecting the particular promoter within this list the user obtains enhancer-gene predictions (see below). Selecting the particular enhancer within this list, the user obtains the H3K27ac, open chromatin, CTCF and cofactor fold-changes and activity levels for this enhancer in every tissue, complemented with a list of genes predicted to be regulated by this enhancer, and information about supported eQTL, GWAS and ChIP-Atlas data.

The “ENHANCER-GENE PAIRS” tab reports specifically EPIraction predictions in a tissue-specific manner. The user provides a gene identifier. The portal reports candidate EPIraction enhancer-gene predictions across all tissues. If the prediction is supported by eQTL data or the enhancer overlaps with a GWAS variant, the portal reports the associated data, tissues and traits. If enhancer overlaps with a ChIP-Seq peak the summary of binding TFs is reported.

---

**Extended Data Fig. 1.**
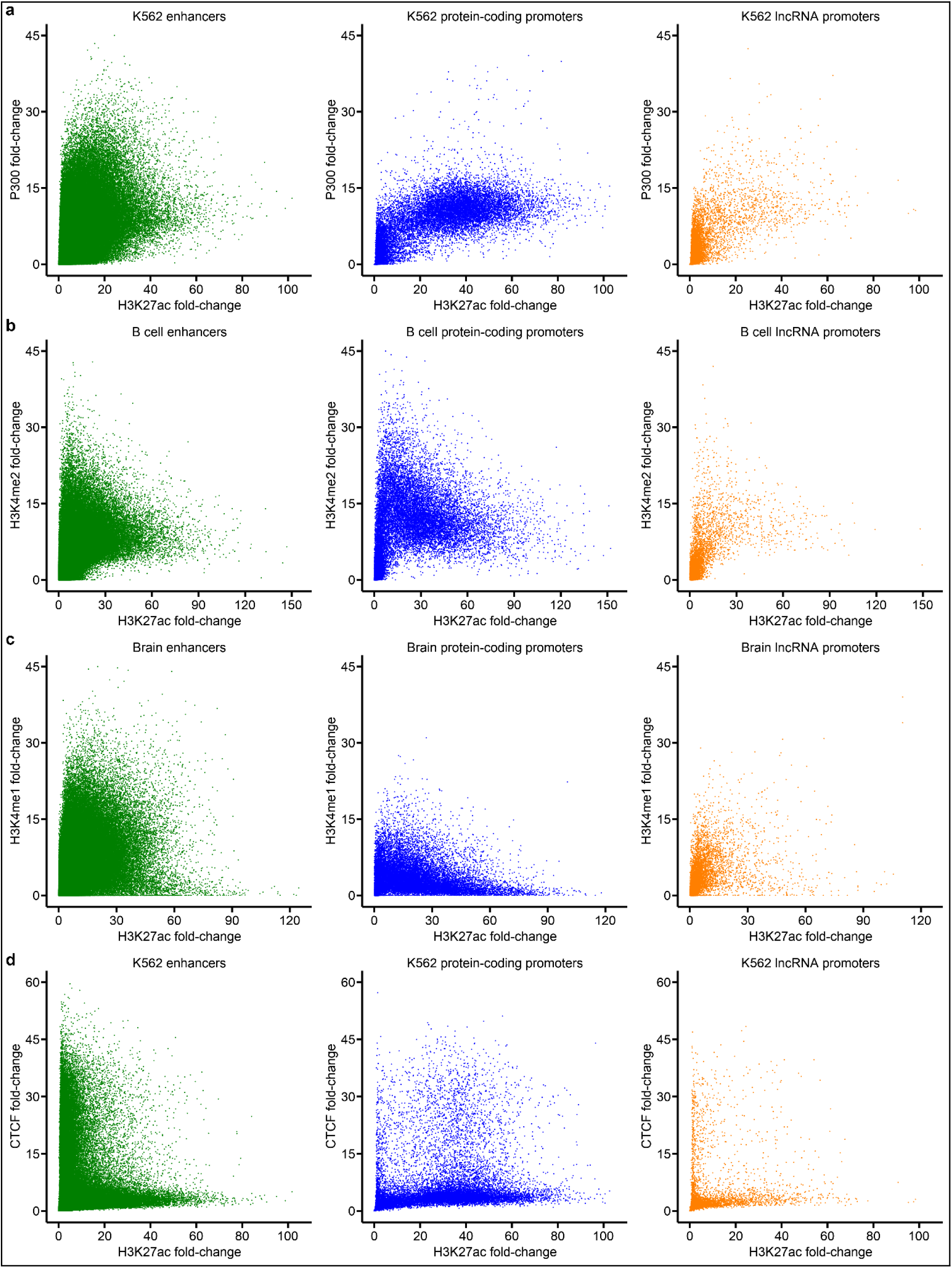
H3K27ac, CTCF and cofactor at enhancer and promoter regions. a-d) Scatterplot of H3K27ac against P300, H3K4me2, H3K4me1 and CTCF signals across candidate enhancers, promoters of protein-coding and lncRNA genes.

**Extended Data Fig. 2.**
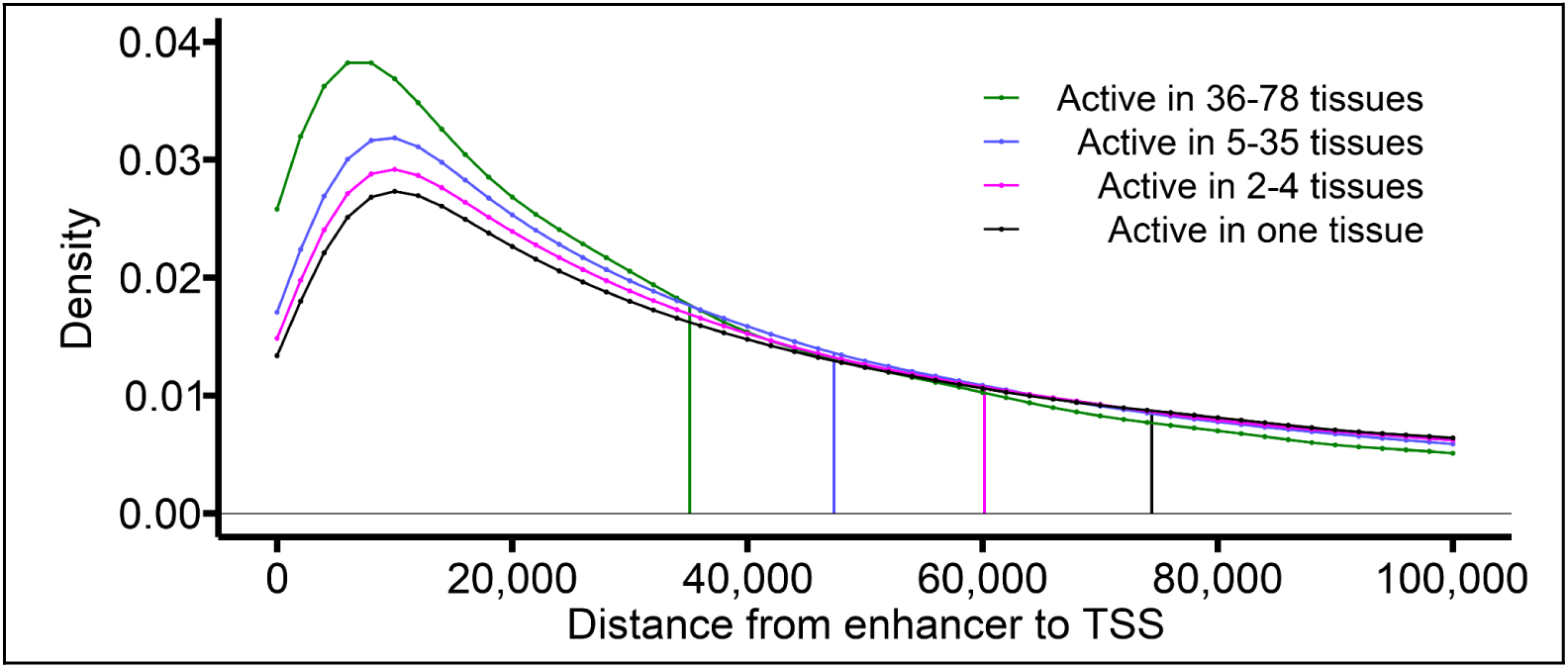
Distribution of the distances from enhancers to the nearest protein-coding gene depending on the number of tissues in which the enhancer is active. The vertical lines show the median values for the corresponding groups.

**Extended Data Fig. 3.**
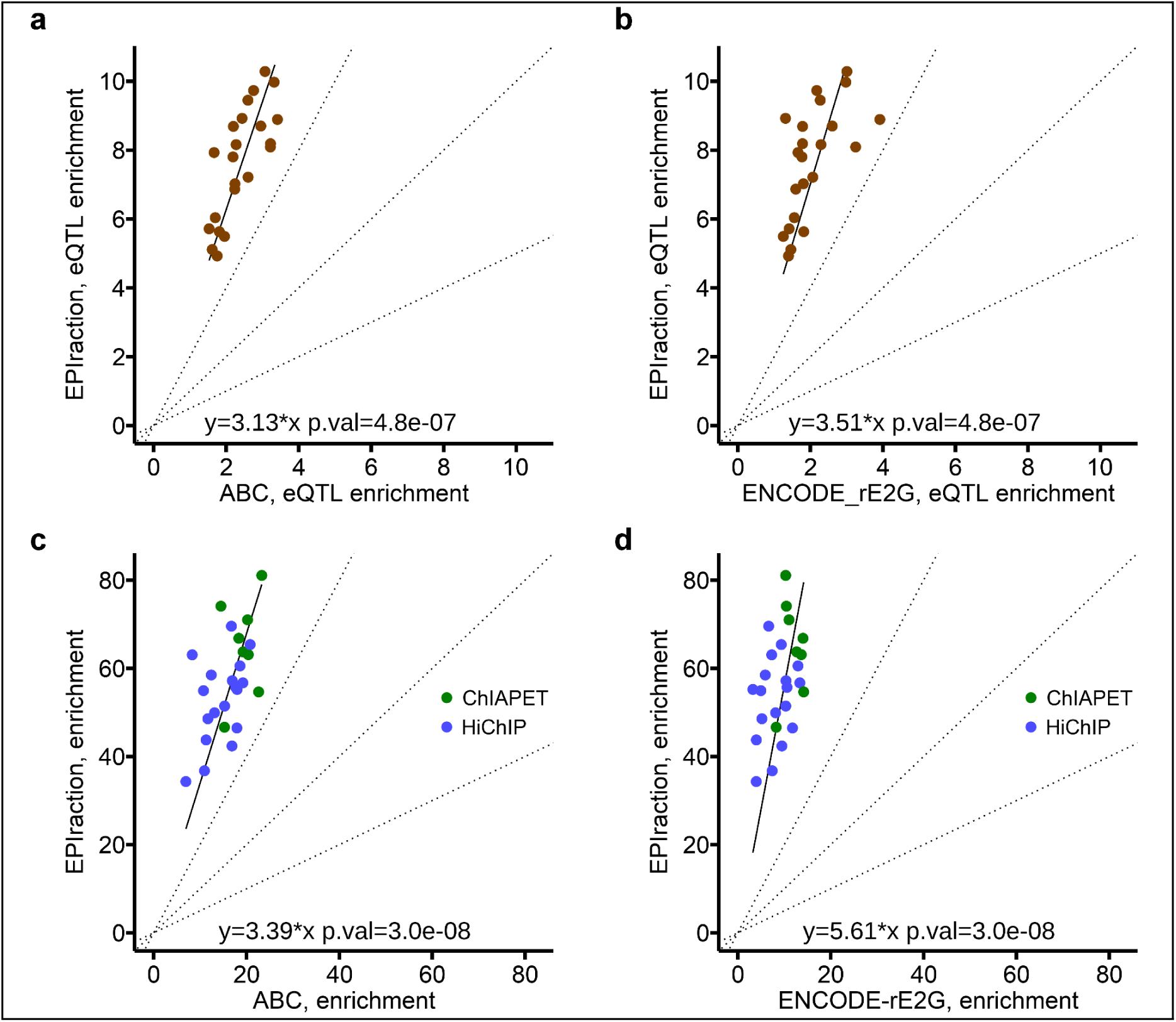
Additional benchmark data of enhancer-gene linking methods. a-b) Comparison of EPIraction, ABC and ENCODE-rE2G predictions at 30% recall on eQTL validation data. P-values are calculated by non-parametric paired Wilcoxon test. c-d) Comparison of EPIraction, ABC and ENCODE-rE2G predictions at 50% recall on HiChIP and ChIA-PET validation data.

**Extended Data Fig. 4.**
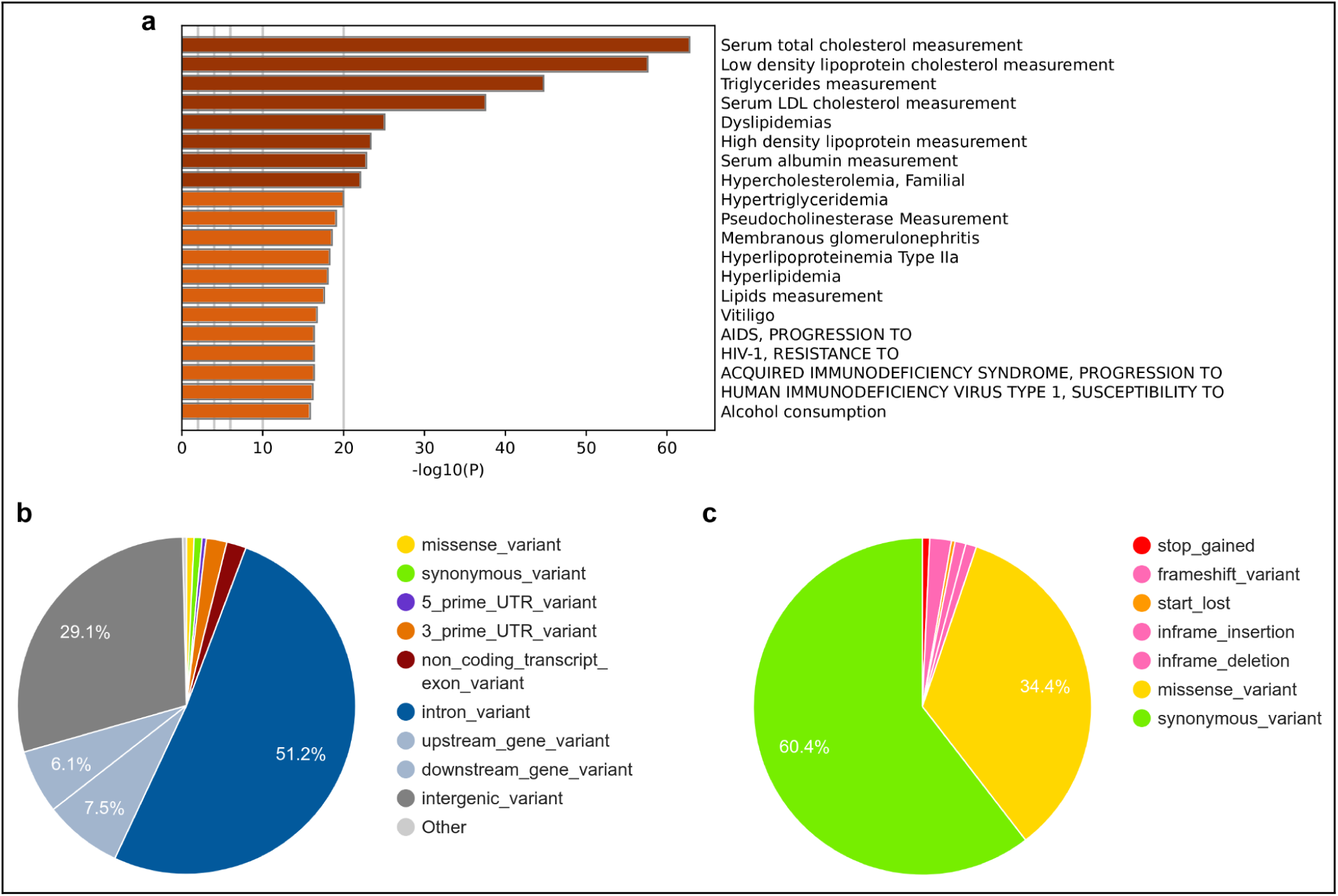
Functional analysis of hypercholesterolemia GWAS SNPs. a) Functional annotation of 867 genes that are predicted to be target genes for hypercholesterolemia SNPs by EPIraction at 0.05 threshold. The DisGeNET (58) annotations are reported. b) Genome-wide annotation of all 12,749 hypercholesterolemia SNPs. c) Annotation of 579 SNPs that lie within the protein-coding regions.

**Extended Data Fig. 5.**
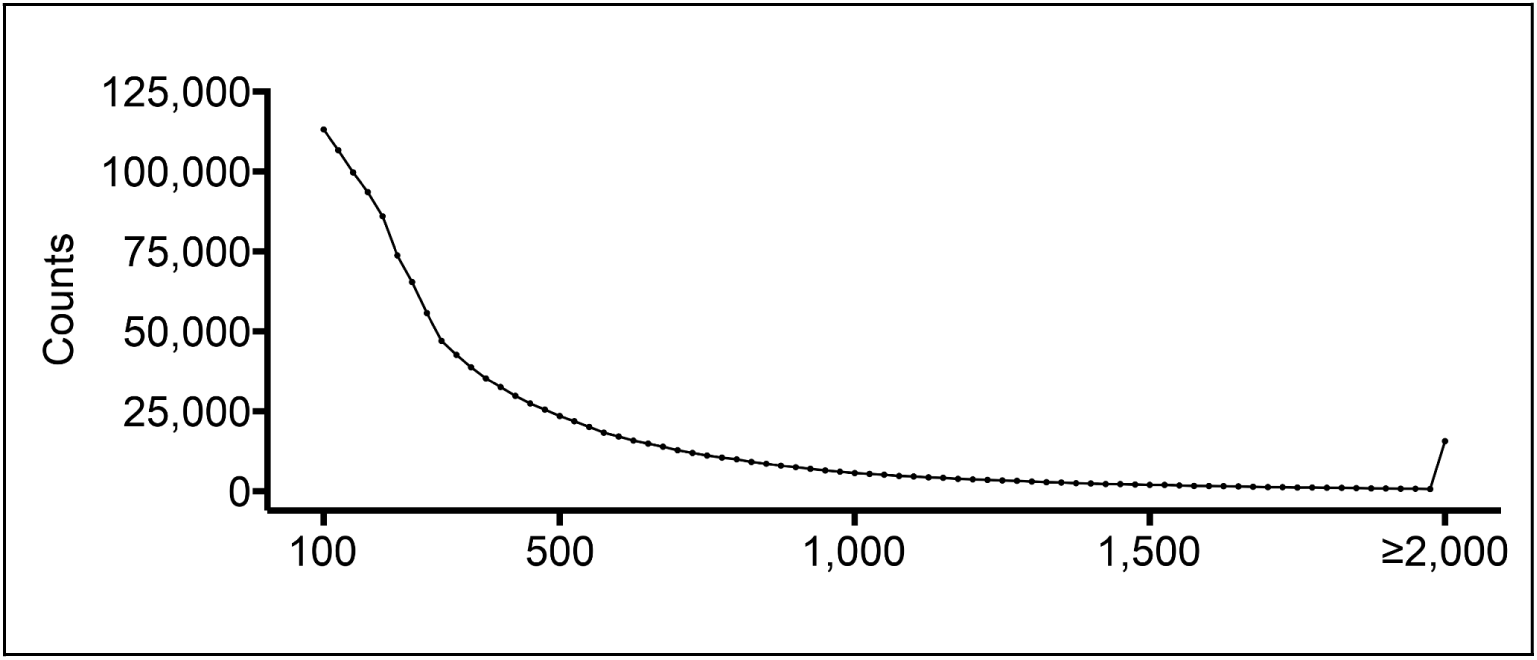
The length distribution of merged open chromatin and CTCF peaks.

**Extended Data Fig. 6.**
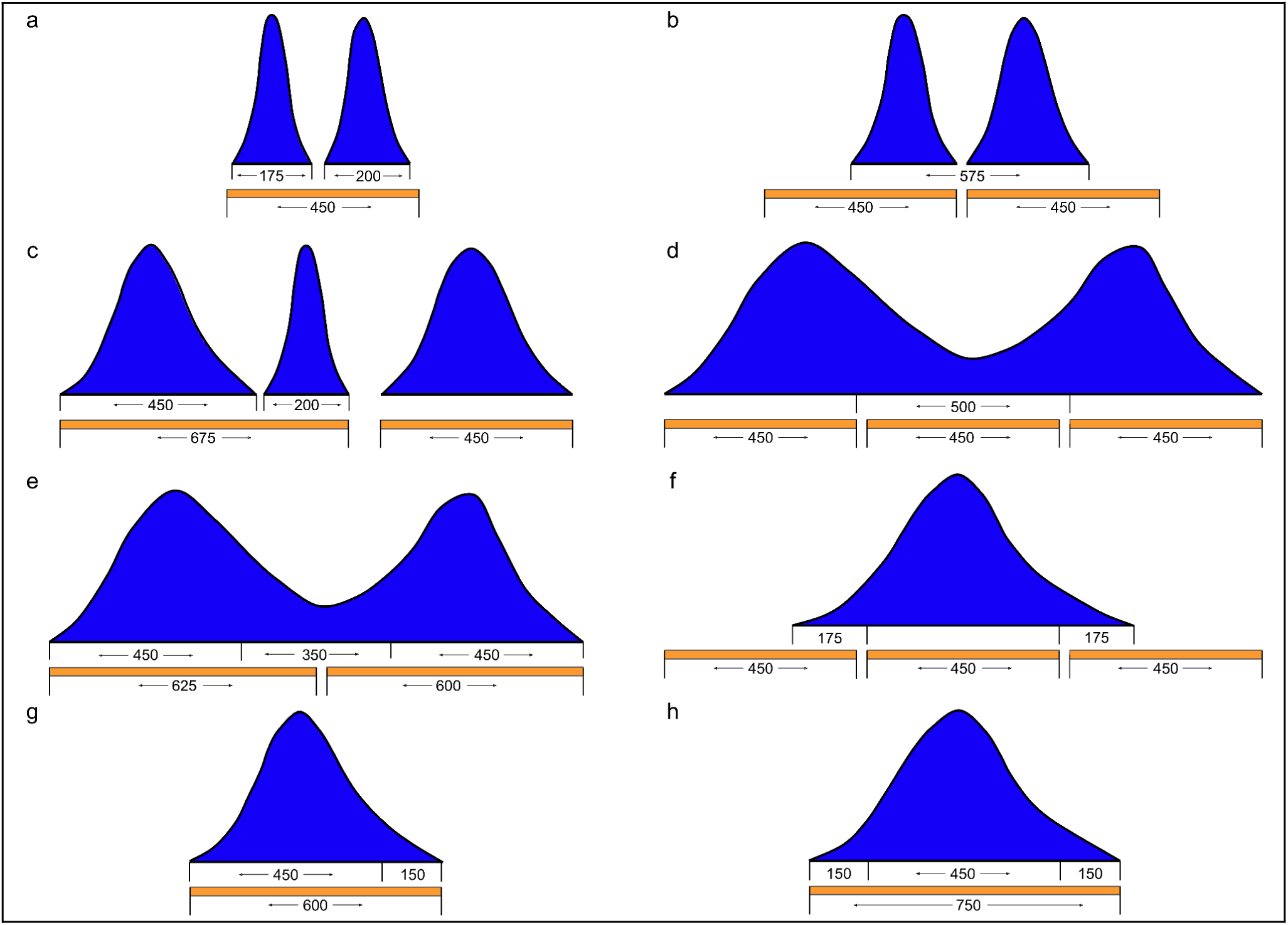
Tissue-agnostic list of candidate enhancers derived from CTCF and open chromatin peaks. Peaks resulting from merging CTCF and chromatin peaks are displayed in blue. The height of the peaks is the sum of the open chromatin and CTCF fold-change of the merged peaks. Candidate enhancer regions are represented by orange rectangles. The numbers indicate the length of regions in base pairs. a) Neighbor short peaks are merged if the length of the merged region does not exceed 450 bp. b) Short peaks are extended to 450 bp in both directions if they do not overlap other peaks. Two neighbor short peaks are extended independently to 450 bp in opposite directions. c) Short peaks that cannot be extended are merged to neighbor peaks, even if the length of the merged peaks is longer than 450 bp. d) 450 bp candidate enhancers are generated within the long peaks. e) 450 bp candidate enhancer regions are extended to close the gap between them, if the extension overlaps some peak. f) Candidate enhancer regions are created up- and/or down-stream of candidate enhancer regions if they overlap peak termini by at least 175 bp. g,h) Candidate enhancer regions are extended up- and/or down-stream if they overlap peak termini insufficiently.

**Extended Data Fig. 7.**
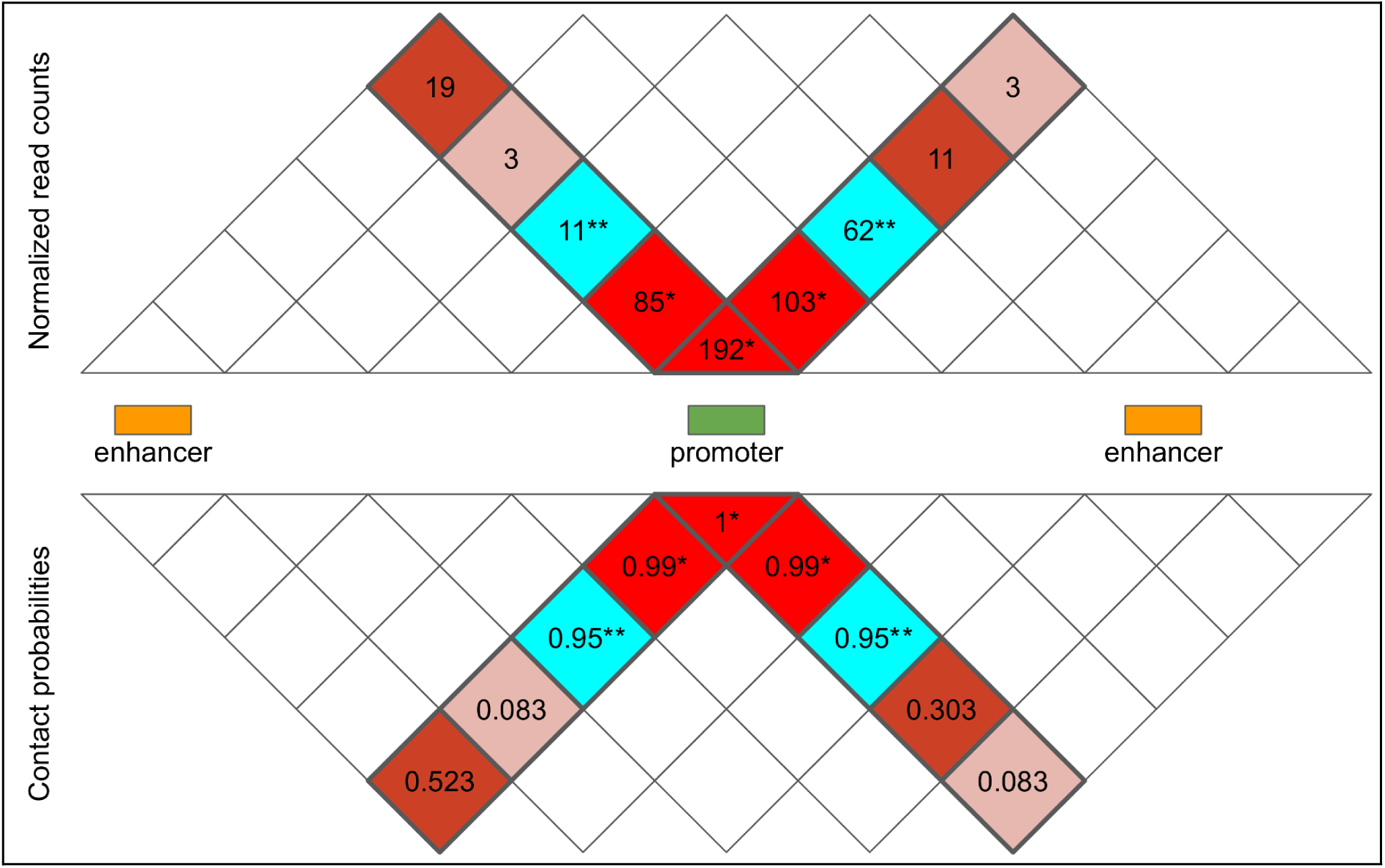
Probability of enhancer-promoter interaction (contact) from normalized Hi-C read count matrix. The zero- and first-diagonal read counts (*) do not participate in calculations. Contact probability on the zero diagonal is set to 1, on the two first diagonals set to 0.99, on the two second diagonals to 0.95. The average of two normalized second-diagonal read counts (**) forms the normalization factor M_p_. Contact probability at third and higher diagonals is set to the number of Hi-C reads supporting this diagonal divided by M_p_ and constrained to the value of 1. The actual read counts are fractional due to normalization. For simplicity they are displayed as integers.

